# Mapping Environmental Discourses: A Bibliometric and Critical Analysis of Eco-Discourses in Scopus

**DOI:** 10.1101/2025.06.07.658453

**Authors:** Martina Paul Chaligha

**Author notes:** **Corresponding Author:** Dr. Martina Paul Chaligha.

## Abstract

This study offers a bibliometric meta-analysis of eco-discourse and discourse analysis within Scopus-indexed journals from 2010 to 2020, focusing on how language frames environmental challenges in global scholarly conversations. Using a systematic PRISMA-based approach, the study integrates bibliometric tools (VOSviewer and Scopus analysis) to trace publication trends, key countries, influential sources, and frequently cited references. Keyword co-occurrence analysis highlights central themes such as discourse analysis, sustainability, ecology, climate change, and Critical Discourse Analysis (CDA) illuminating the evolving intersections of these topics across disciplines. Crucially, this study foregrounds how discourse operates as a social practice in shaping perceptions and actions around ecological crises, revealing ideological framings and power dynamics within environmental discourses. Co-citation analysis further elucidates the field’s theoretical and methodological convergences with sociolinguistics, media studies, and environmental policy discourse. By synthesizing these diverse threads, the study advances critical reflections on the role of discourse in mediating environmental knowledge and action, offering insights relevant for scholars in discourse studies, environmental communication, and beyond.

## Introduction

In recent decades, global environmental difficulties, including climate change, biodiversity loss, and resource depletion, have heightened the necessity for effective communication regarding ecological issues (Crichton, 2018). This has led to the emergence of eco-discourse, which includes the language and communication tactics employed to express environmental issues, promote sustainability, and affect public and policy-related actions (Eliasson, 2015). Since eco-discourse significantly influences societal consciousness and fosters ecological activism, analyzing its patterns, themes, and evolution has become a crucial academic research domain. Discourse analysis, a fundamental aspect of linguistics and communication studies, provides essential tools for investigating how language influences and propagates environmental values (Alexander & Stibbe, 2014). It allows scholars to analyze the narratives, terminologies, and rhetorical methods used in environmental communication, revealing the underlying ideologies, power dynamics, and cultural settings(Wu, 2018). Despite its significance, the domain of eco-discourse remains fragmented, with numerous approaches and techniques needing a cohesive framework. This fragmentation highlights the significance of bibliometric analysis for mapping the intellectual environment, monitoring academic trends, and pinpointing gaps in the current literature. Bibliometric analysis offers a methodical framework for assessing research productivity, impact, and cooperation within a particular discipline (Ellegaard & Wallin, 2015). This study aims to provide a thorough overview of the scientific landscape of eco-discourse and discourse analysis by examining Scopus-indexed journals, an extensive and high-quality database of academic publications. The bibliometric study aims to give academics a thorough overview of eco-discourse research, guiding future inquiries and interdisciplinary cooperation. This work carefully synthesizes the intellectual evolution of the area, enhancing comprehension of the progression of eco-discourse and its ongoing influence on global environmental narratives.

### Literature review

Research in eco-discourse examines the language and communication techniques employed to formulate and propagate environmental narratives, mirroring society’s values, ideologies, and ecological issues. Several studies have delved into Eco-Discourse, analyzing its role in environmental communication and advocacy (Guan-qun, 2023; Richard, 2006; Sean, 2015; Wu, 2018; B. Zhang et al., 2023; W. Zhang & Xiao, 2023; Zollo, 2024). Theoretical frameworks, such as ecolinguistics(Penz & Fill, 2022), Stibbe (2015), and critical discourse analysis (CDA) (Dijk, 1995), underpin this area, highlighting the influence of language on views of environmental challenges. Explored how language and narratives in Eco-Discourse shape public perceptions of environmental issues, focusing on using metaphors and framing devices to evoke emotional and cognitive responses. This foundational work emphasized the persuasive power of Eco-Discourse in fostering environmental awareness. Similarly, Dryzek (1997) examined the role of discourse in environmental policymaking, identifying dominant narrative structures that influence sustainability initiatives. This study underscored the ideological underpinnings of Eco-Discourse and its impact on environmental governance. Another significant contribution is Stibbe’s (2014) “ecolinguistics” analysis, which integrates ecological concerns with linguistic studies, highlighting how language choices reflect and reinforce ecological values. Lakoff (2010) investigated how framing techniques in Eco-Discourse, particularly in climate change communication, shape public and political engagement. Ainsworth (2021) explored how Eco-Discourse operates within educational settings, advocating for integrating ecological narratives in curricula to promote environmental literacy. Collectively, these studies provide a robust theoretical foundation for understanding the mechanisms and impacts of Eco-Discourse in various contexts.

Recent studies highlight Indigenous viewpoints and ethical considerations, promoting eco-centric narratives over anthropocentric ones. Researchers such as Whyte (2018) and Hatley (Hatley, 2016) have examined Indigenous ecological knowledge and its incorporation into eco-centric narratives. Plumwood (2005) challenges anthropocentrism and proposes a transition to eco-centric ethical frameworks. Berkes (2017) underscores the significance of traditional knowledge in sustainable, environmentally friendly behavior.

Although ecological discourse and discourse analysis have gained increasing attention in recent years, comprehensive bibliometric reviews that incorporate PRISMA guidelines alongside VOSviewer software remain limited. This study aims to bridge this gap by systematically reviewing literature related to ecological and environmental discourse, drawing from the Scopus database. The objective is to map research trends, analyze key themes, and provide a structured bibliometric overview of the field. Several factors justify the need for a bibliometric study integrating PRISMA methodology and VOSviewer visualizations. First, a bibliometric analysis not only organizes and synthesizes existing research but also provides critical insights into emerging topics and influential works in the field. Second, identifying foundational studies can be particularly challenging for students and early-career researchers. VOSviewer aids in visualizing citation networks, keyword co-occurrence patterns, and thematic clusters, making it easier to navigate the existing literature. Moreover, PRISMA guidelines enhance the transparency and rigor of systematic literature reviews by offering a structured approach to selecting and analyzing studies. Originally introduced by Moher et al. (2009), PRISMA has become a widely accepted standard for conducting systematic reviews and meta-analyses, ensuring clarity and reproducibility in academic research. In summary, qualitative methodologies predominate, with a growing use of corpus linguistics and bibliometric analysis to discern patterns and knowledge deficiencies. Media studies indicate sensationalism in environmental reporting, although corporate speech frequently underscores discrepancies between sustainability language and actual actions (Lusagalika, 2020). Eco-discourse studies require enhanced incorporation of quantitative methodologies, global views, and multidisciplinary cooperation to tackle intricate environmental issues comprehensively.

## RESEARCH DESIGN

### Methodology

Systematic literature reviews often adhere to the PRISMA guidelines, but recent research has increasingly incorporated analytical software and visualization tools to enhance the synthesis of findings for example (Kalibatiene & Miliauskaitė, 2021; Zupic & Čater, 2015). In particular, Almasri et al. (2021) proposed an adaptation of PRISMA that integrates bibliometric techniques within the standard workflow. Building on this approach, the present study employs a structured methodology that combines PRISMA guidelines with bibliometric analysis, utilizing VOSviewer software as a primary tool. The methodology follows a four-stage process research design, data collection, data analysis, and visualization similar to prior bibliometric studies. Specifically, this research seeks to answer key questions related to ecological discourse and discourse analysis over the period from 2010 to 2020. These include identifying the most active contributing countries, determining the most influential sources, uncovering dominant research themes, and mapping co-citation networks in this domain. By applying bibliometric techniques to data retrieved from Scopus, this study provides a structured and systematic examination of trends in ecological discourse and discourse analysis.

The second stage of the research involves data collection, structured according to the PRISMA approach. Figure 2 presents a modified PRISMA flowchart that maps the selection process of relevant records. Following standard systematic review practices, the data collection phase consists of four main steps: identification, screening, eligibility assessment, and final inclusion. To ensure a rigorous selection process, predefined inclusion and exclusion criteria were established before retrieving the data. The inclusion criteria were set as follows: Studies published in peer-reviewed journals indexed in the Scopus database. Use of key terms related to ecological discourse, including but not limited to “environmental communication,” “ecolinguistics,” “environmental discourse,” and “environmental policy.” Only scholarly articles were considered. The study period covered publications from January 1, 2010, to January 1, 2020. The exclusion criteria were applied to filter out irrelevant or low-quality sources, including: Publications where the terms “ecological discourse” or “discourse analysis” were mentioned without being central to the study. Articles with fewer than five pages. Studies that did not explicitly focus on ecological discourse as a research objective. In the identification phase, relevant articles were retrieved from the Scopus database using a structured search strategy. The selected keywords included “ecological discourse” OR “environmental discourse” OR “environmental communication” OR “environmental policy” OR “ecolinguistics” OR “ecology” AND “discourse.” The initial search process yielded a total of 862 records. In the screening phase, the dataset was refined by selecting only journal articles, thereby excluding book reviews, conference proceedings, meeting abstracts, editorials, and other non-scholarly materials. Additionally, to ensure consistency in language, only publications available in English were retained, reducing the dataset to 589 records. The study period covered research published between January 1, 2010, and January 1, 2020.

**Figure 1.**
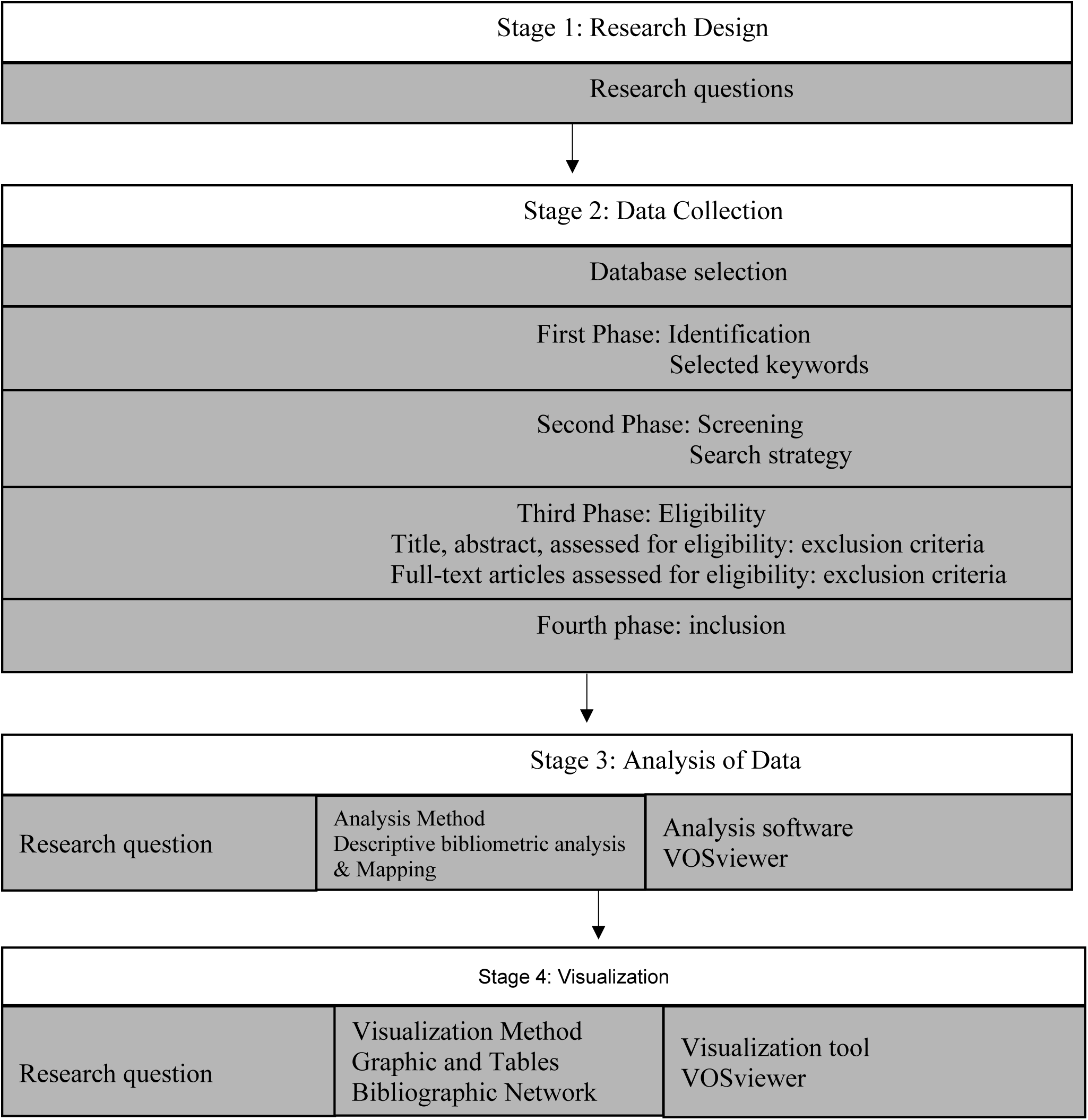
provides a visual representation of the methodological framework, illustrating the integration of PRISMA guidelines with bibliometric tools and visualization software.

**Figure 2.**
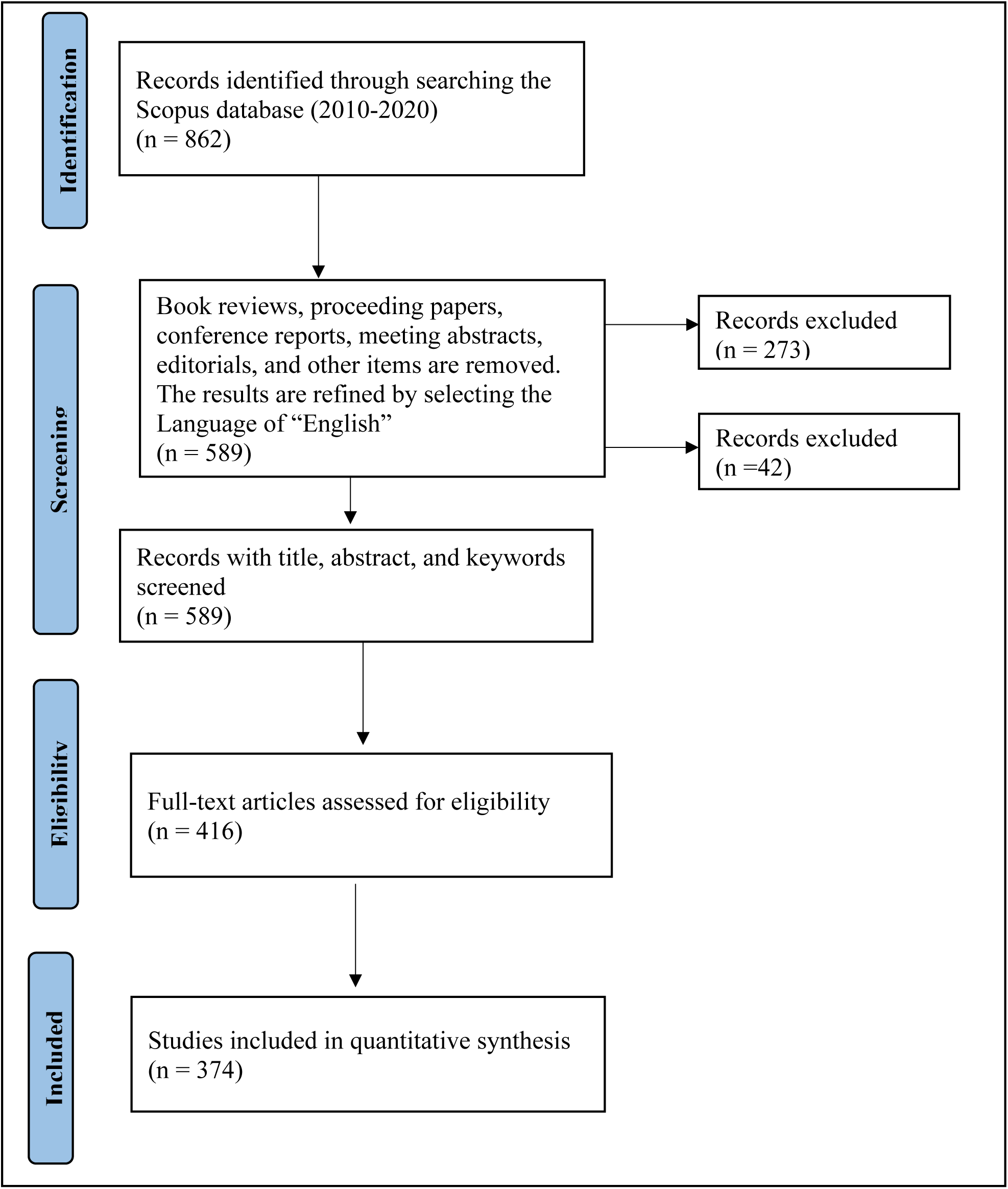
The revised PRISMA flowchart.

During the eligibility assessment, the remaining 416 records underwent a full-text review. This step was necessary to verify whether the identified keywords were central to the study rather than being mentioned in passing or in the background. Articles that did not explicitly focus on ecological discourse and discourse analysis as primary research topics were excluded. As a result, 42 additional records were removed, leading to a final selection of 374 studies for bibliometric analysis.

### Bibliometric Analysis

Following data retrieval, a bibliometric analysis was conducted to examine trends in ecological discourse and discourse analysis within Scopus-indexed literature. The dataset was processed using VOSviewer (van Eck & Waltman, 2010) to perform citation and co-citation analyses. Initially, publication trends and contributions from various countries were visualized using Microsoft Excel based on Scopus data. To address the second research question, citation analysis was conducted using VOSviewer’s citation function, which measures the interconnection between sources based on the frequency of citations among authors. This method provides insight into the academic influence of articles and journals, where higher citation counts indicate greater impact in the field. For the third research question, a network analysis of keyword co-occurrence was performed using Scopus data (Gupta, 2023; Kamarullah, K., & Yanti, 2024). The degree of relatedness between terms was determined based on their co-occurrence frequency, revealing thematic clusters within the literature. To address the fourth research question, a multi-level network and cluster analysis were conducted using VOSviewer. First, co-citation analysis identified the most frequently cited references in the field and their relationships with other works. Second, co-citation patterns among highly referenced sources were examined. Finally, an author-based co-citation analysis was performed to determine the most influential scholars contributing to this research domain. Co-citation analysis quantifies the frequency with which two documents are cited together, reflecting their thematic proximity. This study employed the complete counting method, where all co-occurrence and co-citation links were assigned equal weight, ensuring a comprehensive assessment of citation structures.

## Results and Discussion

**Figure 3** shows that the annual publication on ecological discourse and discourse analysis has been and is still increasing. We selected the year range 2010-2020 for the following reasons: the emergence and growth of eco-discourse research, the advancements in bibliometric tools, the global policy and environmental awareness, and the growth of interdisciplinary research. Between 2010 and 2012, the number of published articles ranged from sixteen to nineteen, indicating that research interest in this topic was relatively limited during that period. From 2013 to 2017, over thirty papers were published, suggesting a growing scholarly interest and momentum in ecological discourse and discourse analysis. From 2017 to 2020, over fifty papers were published suggesting scholars increasingly recognized the importance of analyzing how language constructs ecological issues and shapes environmental narratives. The rapid expansion reflects a response to pressing global ecological crises such as climate change.

**Figure 3.**
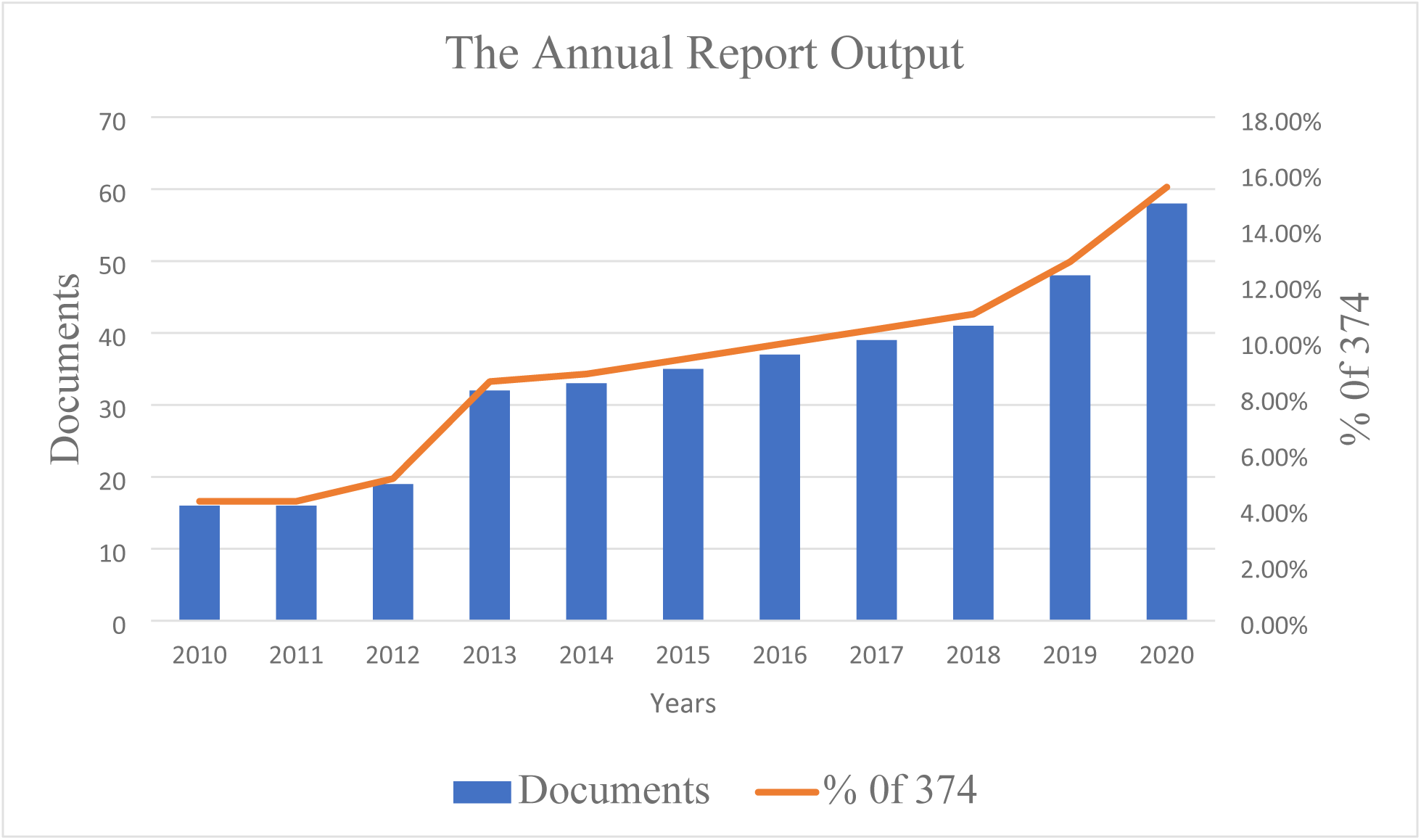
Trends in Annual Research Publications

**Table 1** presents the top ten most active countries in ecological discourse research. Out of 62 countries, only those with at least ten published papers were included in this analysis. The second column lists these countries, while the third and fourth columns display the number of publications and their corresponding percentage of the total 374 articles. Researchers from the United States contributed the highest number of publications between 2010 and 2020, accounting for 24.33% of the total output. Other leading contributors include the United Kingdom (14.70%), Germany (7.75%), Australia (7.48%), Canada (6.95%), Sweden (5.88%), the Netherlands (4.01%), China (3.74%), and Belgium (2.94%). In contrast, Spain produced fewer than ten papers, indicating relatively lower research activity in this area.

**Table 1.**
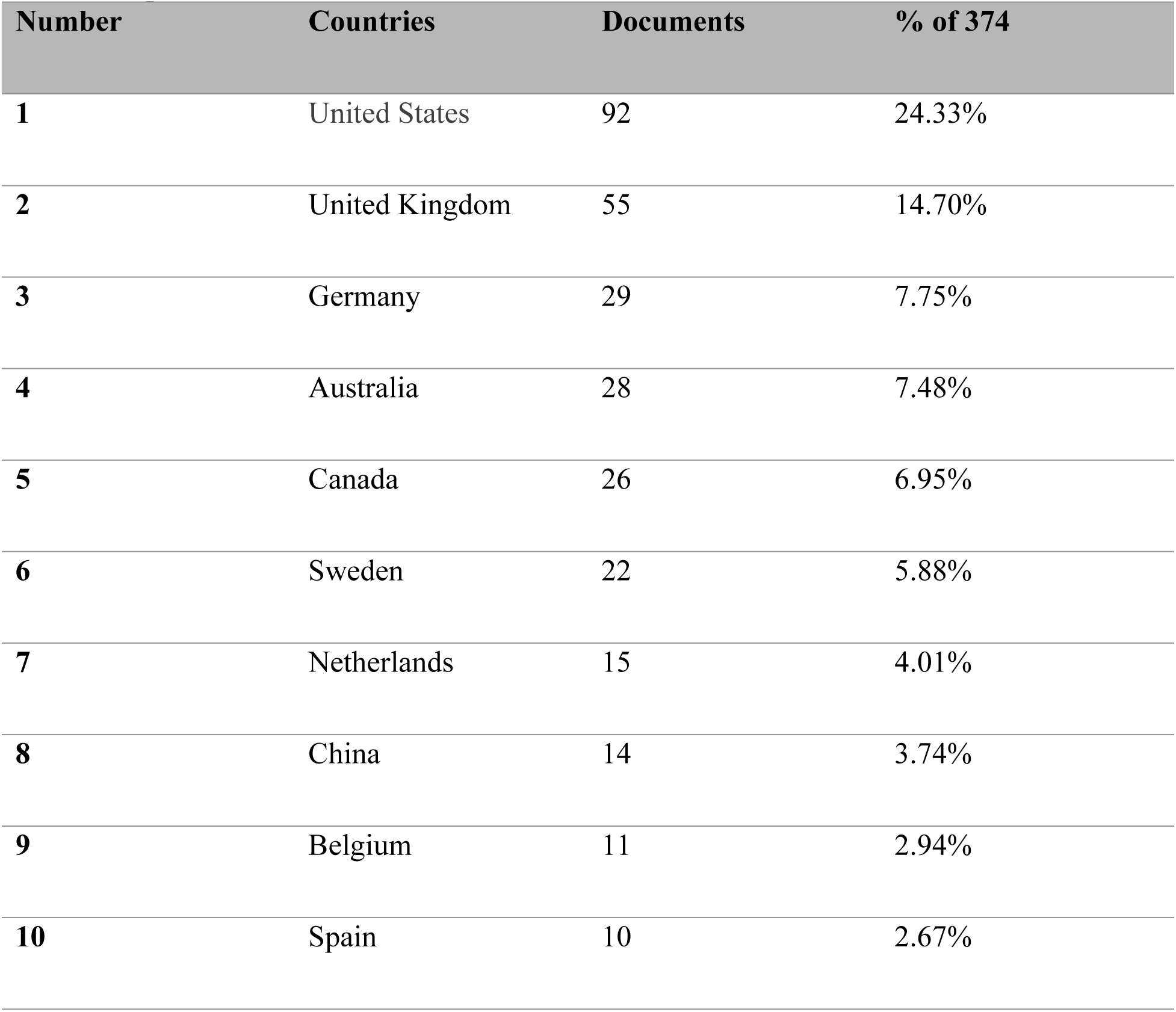
Top Ten Productive Countries/Territories.

A total of 374 articles were distributed across 310 different journals. To evaluate the significance of various journals in ecological discourse research, a threshold was established requiring at least two published documents per source and a minimum of four citations. Of the 310 journals reviewed, Table 2 presents the ten most frequently cited. The second and third columns in the table indicate the number of published articles and their corresponding citation counts. Among these, Ecology and Society stands out as a leading journal, publishing ten articles and achieving a high citation frequency, which underscores its influence in this research area. A citation analysis using VOSviewer identified Gregson’s (2015) article, “Interrogating the Circular Economy: The Moral Economy of Resource Recovery in the EU”, as the most cited work in this journal, with 582 citations. Furthermore, Journal of Environmental Policy and Planning, Geoforum, and Ecological Economics have each received over 300 citations in Scopus, with more than five articles published in each journal. Similarly, Global Environmental Change contains five articles, while Sustainability (Switzerland) has 13, both surpassing 200 citations. Additional journals such as Environmental Politics, Critical Discourse Studies, Biodiversity and Conservation, and Discourse and Communication have published a smaller number of articles, ranging from one to six. The overall impact of a journal is determined by considering both its citation count and the volume of research it has contributed to the field(Huang et al., 2024).

**Table 2.** Top Ten Sources.

| **Journal titles** | **Sum of Citations** | **Sum of No. of articles** |
| --- | --- | --- |
| 1. Ecology and society | 582 | 10 |
| 2. Journal of Environmental Policy and Planning | 369 | 6 |
| 3. Geoforum | 334 | 7 |
| 4. Ecological economics | 307 | 10 |
| 5. Global environmental Change | 279 | 5 |
| 6. Sustainability (Switzerland) | 239 | 13 |
| 7. Environmental Politics | 192 | 5 |
| 8. Critical discourse studies | 86 | 2 |
| 9. Biodiversity and conservation | 70 | 6 |
| 10. Discourse and Communication | 22 | 1 |

### Co-occurrence Analysis of Keywords

Keyword searches were conducted based on the terms provided by the authors of the 374 papers. After consolidating similar keywords, a total of 1,419 distinct terms were identified. To refine the analysis, a minimum occurrence threshold of two was applied, resulting in 180 keywords that met the criteria. Table 3 presents the top ten most frequently occurring keywords. The keyword “Discourse Analysis” has the highest frequency, appearing forty times, followed by “discourse,” which appears twenty-two times. This suggests that scholars view discourse as a fundamental tool for understanding and critiquing language in diverse social, political, and environmental contexts. This supports the findings of McDonald’s (2013) research on the discourses surrounding climate security, which bridges the gap between discourse studies and environmental studies.

**Table 3.** Top Fifteen Keyword List.

| Number | Keywords | Occurrence |
| --- | --- | --- |
| 1. | Discourse Analysis | 40 |
| 2. | Discourse | 22 |
| 3. | Climate Change | 18 |
| 4. | Sustainability | 21 |
| 5. | Ecology | 11 |
| 6. | Ecological Modernization | 13 |
| 7. | Critical Discourse Analysis | 13 |
| 8. | Ecosystem services | 12 |
| 9. | Sustainable development | 10 |
| 10. | Environmentalism | 5 |

VOSviewer applies a clustering technique that groups keywords based on their citation relationships (van Eck & Waltman, 2017). Through this automated process, 23 keywords were categorized into four major clusters, each representing a distinct research theme within ecological discourse and discourse analysis. Table 4, along with Figures 4 and 5, visually depicts these keyword clusters, offering a comprehensive view of their distribution. In the network visualization (Figure 4), each node corresponds to a keyword, with node size reflecting the frequency of keyword occurrences. Additionally, in the density visualization (Figure 5), different colors are used to distinguish clusters, highlighting their thematic connections. The following section provides a detailed examination of the four identified clusters.

**Figure 4.**
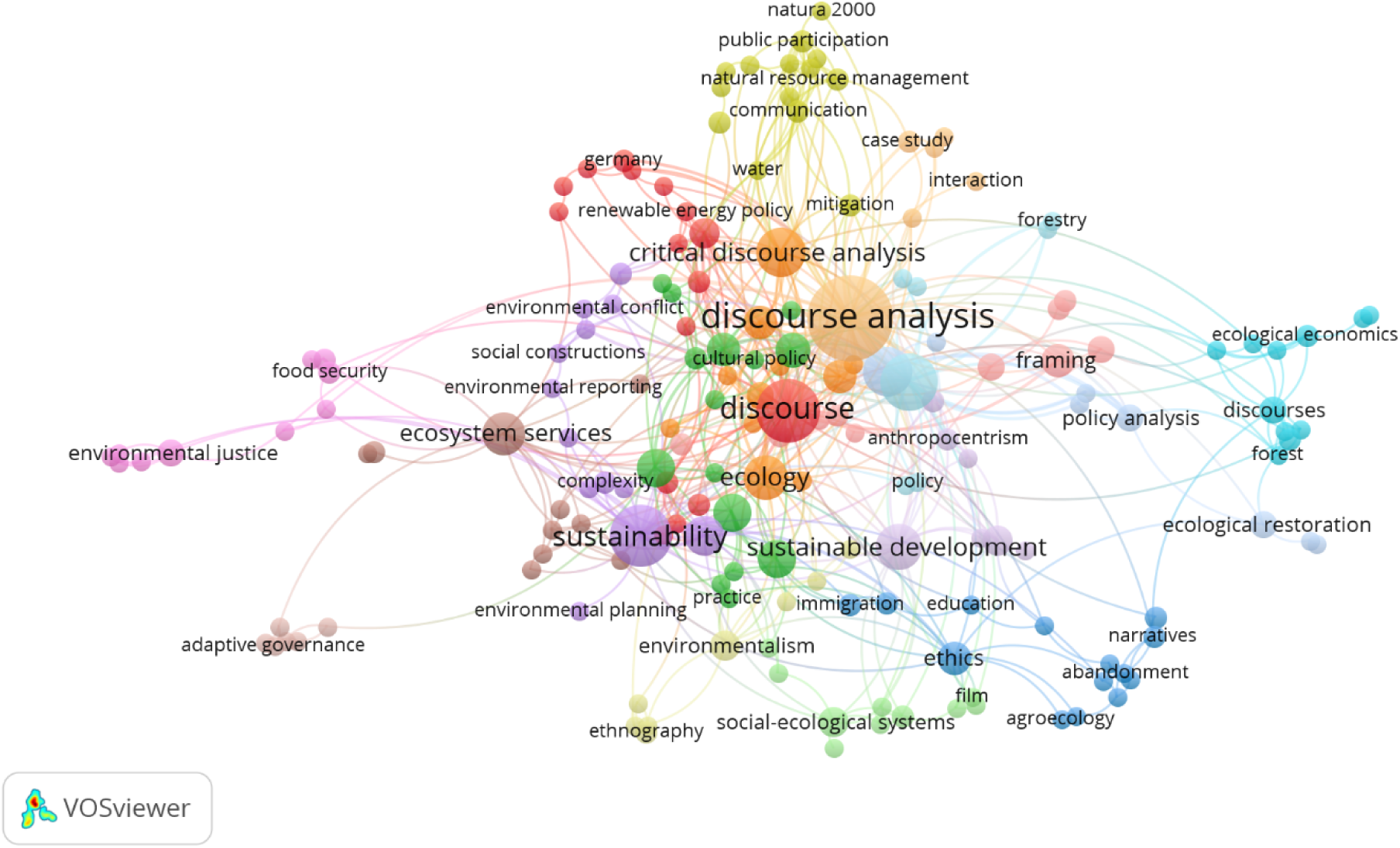
Keyword Network Visualization.

**Figure 5.**
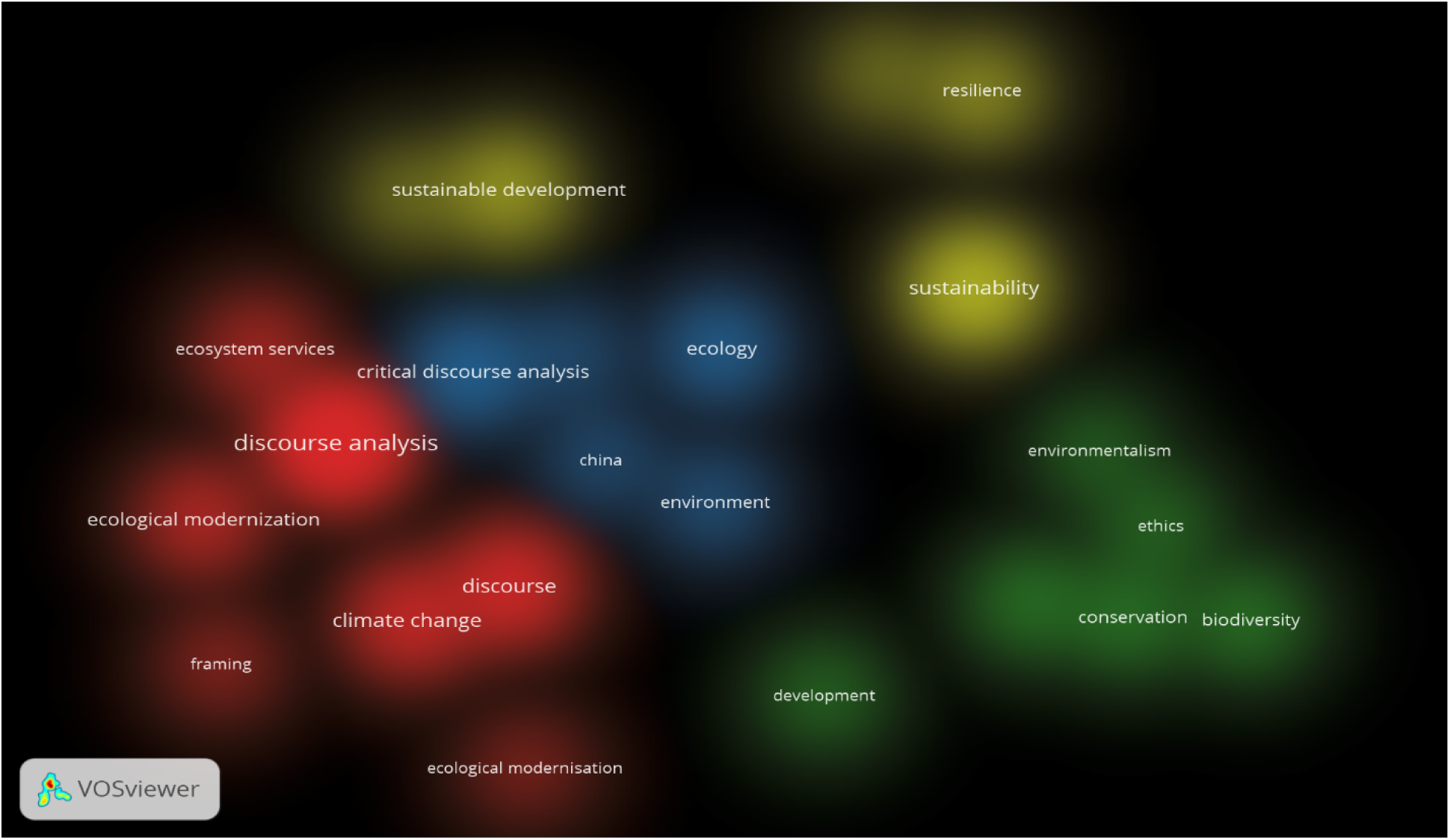
Keyword Density Visualization.

**Table 4.**
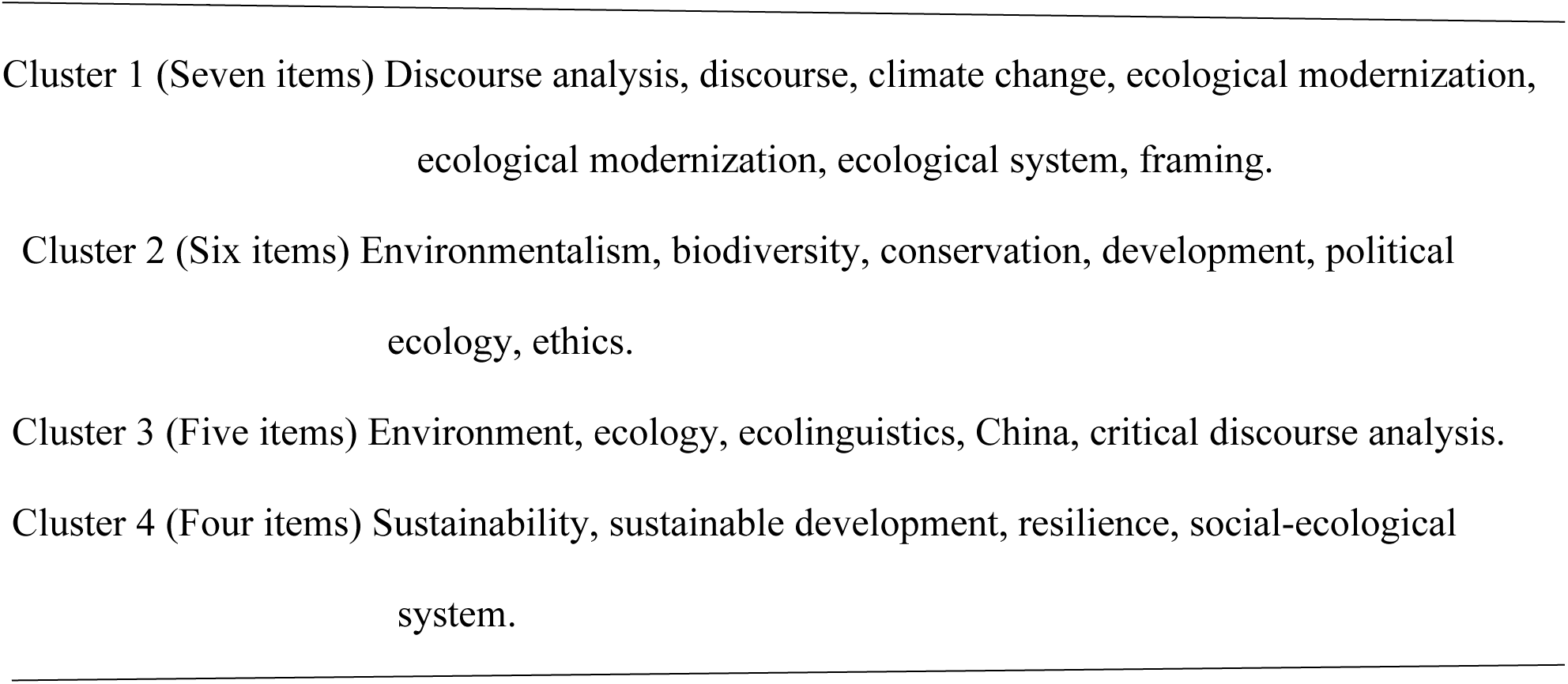
Keyword Clusters.

The red cluster (cluster 1) includes *discourse analysis*, *discourse, climate change, ecological modernization,* and *framing*. *Discourse analysis* suggests that the cluster deals with how language constructs meaning around ecological and environmental issues. Terms such as *climate change* and *framing* underscore the specific emphasis on discussing, framing, and shaping environmental problems in public and media discourse. The research is quantitative (e.g., corpus-based studies) and qualitative methods (e.g., critical discourse analysis), and the research material used in examining ecological discourse and discourse analysis is related to *framing* and *ecological modernization*, as indicated by the term’s ecological modernization and framing. Gong and Philology (Gong & Philology, 2019) adopt a corpus-based discourse analysis to investigate how environmental reports are analyzed from the perspective of ecolinguistics; their findings suggest the promotion of ecological studies. Studies in this area often focus on how language policy discourses across space and time scales (Hult, 2010). Employing critical discourse analysis aims to uncover underlying ideologies and power structures embedded in such narratives, contributing to a broader understanding of environmental communication and its societal impacts (Nguyen, 2023).

The green cluster (cluster 2) contains terms such as *political ecology, conservation*, *biodiversity*, *ethics*, *environmentalism*, and *development.* This cluster focuses on various themes. The field of *political ecology* encompasses environmental ethics, conservation, and biodiversity. It highlights the interaction between social, economic, and environmental factors in shaping ecological outcomes. Research within this cluster often combines qualitative case studies and theoretical frameworks such as political ecology and environmental ethics to investigate conflicts and solutions surrounding conservation and biodiversity (Asiyanbi, 2016; Turner, 2014). Adams (2007) illustrates the integration of political ecology, conservation, biodiversity, ethics, environmentalism, and development, providing valuable insights into the complex relationships between human societies and the natural environment.

Cluster 3, represented in yellow, consists of key terms such as sustainability, sustainable development, resilience, and social-ecological systems. This cluster primarily focuses on sustainability science and resilience theory, highlighting efforts to analyze and strengthen the ability of social-ecological systems to adapt to and recover from environmental challenges. Additionally, the theme of sustainable development links ecological resilience to policy initiatives, including the United Nations Sustainable Development Goals (SDGs). The existence of social-ecological systems indicates the need for integrative approaches that perceive humans and nature as interconnected systems that need systemic solutions. Research in this cluster (García-Rosell & Mäkinen, 2013; van Zeijl-Rozema et al., 2011) often employs quantitative modeling and qualitative assessments to develop frameworks for managing sustainability challenges. It bridges science, governance, and society in achieving resilience to environmental crises.

The blue cluster (cluster 4) features terms such as *ecology, environment, critical discourse analysis, ecolinguistic*s, and *China*. This cluster highlights the role of discourse and language in framing ecological issues, combining linguistic analysis with environmental studies. The keywords *ecology* and *environment* link the linguistic focus to broader ecological studies, examining how language shapes environmental awareness and policy. The keyword China indicates an interest in how China’s environmental practices and policies are discussed and represented globally (Haddad, 2015; Kahn, 2016; Xiong, 2014). Research in this cluster uses corpus-based studies and qualitative discourse analysis to examine environmental communication in political, media, and public domains. It explores the intersection of language, power, and ecological narratives.

### Co-citation Analysis

The analysis began with the co-citation of cited references to determine the most influential works and their interconnections. This approach uses cited references as the primary unit of analysis. As noted by Dong and Chen (2015), the significance of a reference is assessed based on the number of co-citations it may possess. A minimum citation threshold of four was applied, resulting in 45 references meeting this criterion out of a total of 22,244 cited references. VOSviewer was then used to compute the co-citation links among these references, mapping their relationships within the field.

As shown in Table 5, the majority of the 45 cited references consist of books and book chapters, with the exception of ten journal articles (highlighted in bold). These references primarily explore topics such as discourse analysis, environmental discourse, political ecology, governance of ecological systems, and cultural critiques of nature. They provide theoretical, critical, and historical insights into the connections between society, language, and the environment. For example, Fairclough’s (1992)work, which has been cited eight times, is widely recognized for its theoretical and methodological contributions to language and discourse analysis, particularly in examining the relationships between discourse, power, ideology, and social structures. Using VOSviewer, the 45 cited references were categorized into seven distinct clusters. In Figure 6, each node represents a reference, labeled with the author’s name and title. The size of the node reflects its citation frequency, with larger nodes indicating higher citation counts. A connecting line between two nodes signifies that the references have been co-cited at least once. In the co-citation density visualization (Figure 7), different colors distinguish the various clusters, illustrating the thematic connections among the references.

**Figure 6.**
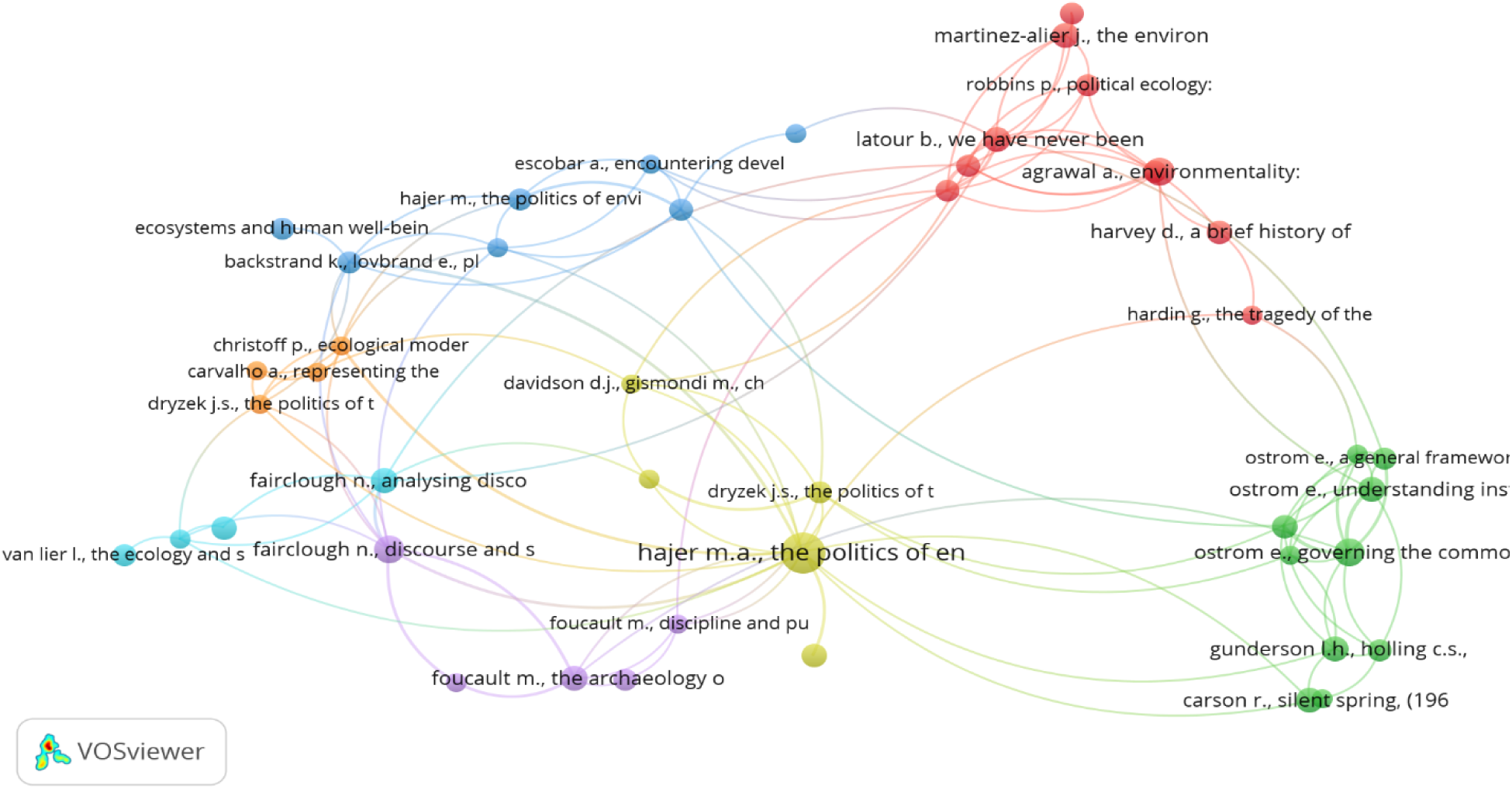
Network Visualization of Co-Citation of The Selected Forty-Five Cited References

**Figure 7.**
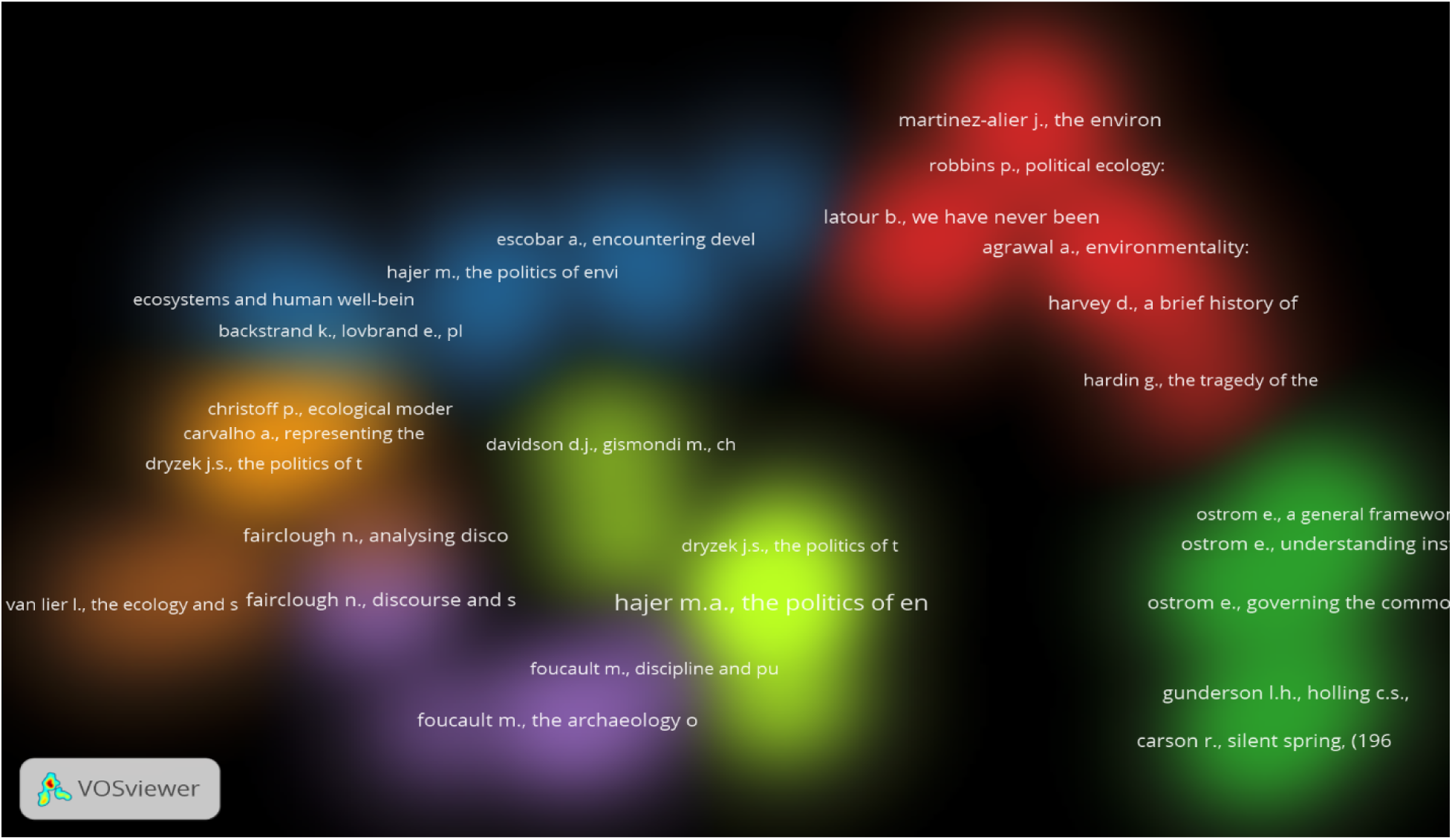
Density Visualization of The Co-Citation of The Forty-Five Cited References

**Table 5.** A list of 45 cited references along with their citation counts and co-citation connections.

| No | Cited references | Citations | Co-citation link |
| --- | --- | --- | --- |
| 1 | Agrawal (2005), environmentality | 8 | 5 |
| 2 | **Backstrand, lovbrand (2006), planting trees to mitigate climate change: v 6, p. 50-75** | 5 | 9 |
| 3 | Bennett (2010), vibrant matter: a political ecology of things | 5 | 12 |
| 4 | Blommaert(2005) , discourse: a critical introduction | 4 | 6 |
| 5 | Carson(1962) , silent spring | 7 | 3 |
| 6 | **Carvalho (2005), representing the politics of the greenhouse effect: critical discourse studies, v 2, p. 1-29** | 4 | 4 |
| 7 | **Christoff (1996), ecological modernization, ecological modernities, environmental politics, v 5, p. 476-500** | 4 | 9 |
| 8 | Cronon (1996), the trouble with wilderness | 5 | 5 |
| 9 | Davidson, gismondi (2011), challenging legitimacy at the precipice of energy calamity | 4 | 6 |
| 10 | Dryzek (2005), the politics of the earth: environmental discourses | 4 | 7 |
| 11 | Dryzek (2013), the politics of the earth: environmental discourses | 5 | 8 |
| 12 | Millennium (2005), ecosystems and human well-being: Synthesis | 5 | 1 |
| 13 | Escobar (1995), encountering development | 4 | 5 |
| 14 | Fairclough (2003), analyzing discourse: textual analysis for social research | 7 | 7 |
| 15 | Fairclough and Wodak (1997), Discourse as Social Interaction | 4 | 4 |
| 16 | Fairclough (1995), critical discourse analysis: the critical study of language | 4 | 7 |
| 17 | Fairclough (1992), discourse and social change | 8 | 13 |
| 18 | Fairclough (2001), language and power | 4 | 5 |
| 19 | **Folke, hahn, olsson, norberg (2005), adaptive governance of social-ecological systems, v 30, p.441-473** | 6 | 10 |
| 20 | Foucault (1977), discipline and punish: the birth of the prison | 4 | 5 |
| 21 | Foucault (1972), the archaeology of knowledge | 7 | 8 |
| 22 | **Garcia, martin, gonzalez, montes (2008), social perceptions of the impacts and benefits of invasive alien species, v 12, p. 2969-2983** | 4 | 2 |
| 23 | Goffman (1974), frame analysis: an essay on the organization of experience | 6 | 2 |
| 24 | Gunderson, holling, panarchy (2002), understanding transformations in human and natural systems | 7 | 6 |
| 25 | Hajer (1997), the politics of environmental discourse | 5 | 6 |
| 26 | **Hajer, versteeg (2005), a decade of discourse analysis of environmental politics, v 3, p. 175-184** | 5 | 8 |
| 27 | Hajer (1995), the politics of environmental discourse | 18 | 20 |
| 28 | **Hardin (1968), the tragedy of the commons, science, v 162, p. 1243-1248** | 4 | 3 |
| 29 | Harvey (2005), a brief history of neoliberalism, (2005) | 6 | 3 |
| 30 | **Holling (1973), resilience and stability of ecological systems, v 4, p. 1-23** | 4 | 5 |
| 31 | Hulme (2009), why we disagree about climate change: understanding controversy | 4 | 3 |
| 32 | Jorgensen, phillips (2002), discourse analysis as theory and method | 5 | 2 |
| 33 | Kress, van (1996), reading images: the grammar of visual design | 4 | 3 |
| 34 | Lakoff., johnson (1980), metaphors we live by | 6 | 2 |
| 35 | Latour (2004), politics of nature: how to bring the sciences into democracy | 5 | 11 |
| 36 | Latour (1993), we have never been modern | 7 | 13 |
| 37 | Martinez-alier (2002), the environmentalism of the poor | 7 | 5 |
| 38 | Nixon (2011), slow violence and the environmentalism of the poor | 5 | 2 |
| 39 | **Ostrom (2007), a diagnostic approach for going beyond panaceas, p.181-187** | 4 | 7 |
| 40 | **Ostrom (2009), a general framework for analyzing sustainability of social-ecological systems, v 325, p. 419-422** | 5 | 6 |
| 41 | Ostrom (1990), governing the commons | 8 | 14 |
| 42 | Ostrom(2006) , understanding institutional diversity | 7 | 9 |
| 43 | Robbins (2012), political ecology | 5 | 5 |
| 44 | Sen (1999)development as freedom | 5 | 5 |
| 45 | Van (Van Lier, 2004), the ecology and semiotics of language learning | 5 | 1 |
| Total |  | 249 | 282 |

Cluster 1 (red cluster) primarily concerns political ecology and the broader sociopolitical dynamics of environmental discourse. Key scholars include Martinez-Alier (2002), “The Environmentalism of the Poor,” and Robbins (2012), as well as “Political Ecology,” which address how political and economic systems interact with ecological concerns. Latour (2004), “We Have Never Been Modern” and “Politics of Nature,” explores the intersection of science, politics, and nature. Nixon (2011), in “Slow Violence and the Environmentalism of the Poor,” discusses environmental injustice and the slow, incremental damage caused by ecological crises. This cluster highlights critical approaches to understanding environmental issues, focusing on power relations, environmental justice, and the socio-political framing of ecological problems.

Cluster 2 (the green cluster) centers on ecological systems, governance, and institutional analysis. Ostrom (1990), (“Governing the Commons” and “A General Framework for Analyzing Sustainability”), focused on key studies on resource governance and institutional frameworks for sustainable environmental management. Folke, C., Hahn, T., Olsson, P. (2005), contributions on resilience and adaptive governance in ecological systems. Gunderson, L. H. and Holling, C. S. (2002), foundational studies on ecological resilience and systems theory. This cluster emphasizes the governance of common resources, resilience thinking, and sustainable management frameworks in ecological discourse.

Cluster 3 (Blue Cluster) focuses on developmental discourse, human well-being, and critical discourse analysis. Significant contributions include Escobar (1995) (Encountering Development), which critiques development discourse and its ecological implications. Hajer (1997) (The Politics of Environmental Discourse) explores the role of language and power in shaping environmental policy and discourse. Backstrand, K., and Lovbrand, E. (2006) address ecosystems, sustainability, and human well-being. This cluster highlights critical development studies and the role of discourse in framing environmental and sustainability policies.

Cluster 4 (Yellow Cluster) primarily concerns discourse analysis models and political communication. Key authors such as Fairclough, N (2001) (Language and Power, Analyzing Discourse), foundational works in critical discourse analysis (CDA) that explore language, power, and ideology. Dryzek, J. S. (2005), (The Politics of the Earth), focuses on environmental discourse and its role in political communication. Davidson, D. J., and Gismondi, M. (2011) examine environmental issues’ political and sociological dimensions. This cluster bridges critical discourse analysis and environmental communication, highlighting how political and ecological narratives are constructed.

Cluster 5 (Orange Cluster) addresses media representation of ecological issues and environmental modernity. Christoff, P. (1996), Ecological Modernization, explores how environmental concerns are integrated into modernity and policy. Carvalho, A. (2005), Representing the Environment, focuses on media discourse and the framing of ecological issues. Dryzek, J. S. (2013), connects political discourse with environmental representation. This cluster highlights how media and political discourse shape public perceptions of environmental issues and promote or challenge dominant ecological narratives.

Cluster 6 (Purple Cluster): The purple cluster emphasizes philosophical and ethical dimensions of discourse. Key scholars such as Foucault, M. (1977) (Discipline and Punish, The Archaeology of Knowledge), foundational theories on discourse, power, and knowledge production. Goffman, E. (1974) (Frame Analysis) explores how narratives are framed to shape meaning and perceptions. Lakoff, G., and Johnson, M. (1980) (Metaphors We Live By) highlight the role of metaphor in shaping ecological understanding. This cluster focuses on the philosophical underpinnings of discourse, particularly the ways language frames ethical and cognitive perceptions of ecological issues.

### Co-citation of cited sources

The co-citation analysis of cited sources aims to determine the most frequently co-cited sources and their interconnections. This analysis uses cited sources as the primary unit of study and complements the co-citation analysis of individual references. A minimum citation threshold of 51 was applied, and out of 11,890 sources, 14 met this criterion. These sources are listed in Table 6. As shown in Table 6, Science emerges as the most highly cited and co-cited source, indicating its central role in disseminating influential research across multiple disciplines, including environmental science, policy, and communication. This highlights the journal’s significance in fostering interdisciplinary discussions and advancing scholarship in ecological discourse and related fields. The second most co-cited journal is *Ecology* and *Society*, followed by *Geoforum* in third place. Additional co-citation relationships among these sources are further illustrated in the network visualization (Figure 8). In Figure 8, each node represents a cited source, with labels indicating the journal title. The size of each node corresponds to the number of times the source has been cited, while the proximity between nodes reflects the strength of their co-citation relationships. A closer distance between nodes signifies a stronger thematic or citation-based connection. Figure 9 presents the density visualization, offering further insights into the distribution of co-cited sources

**Figure 8.**
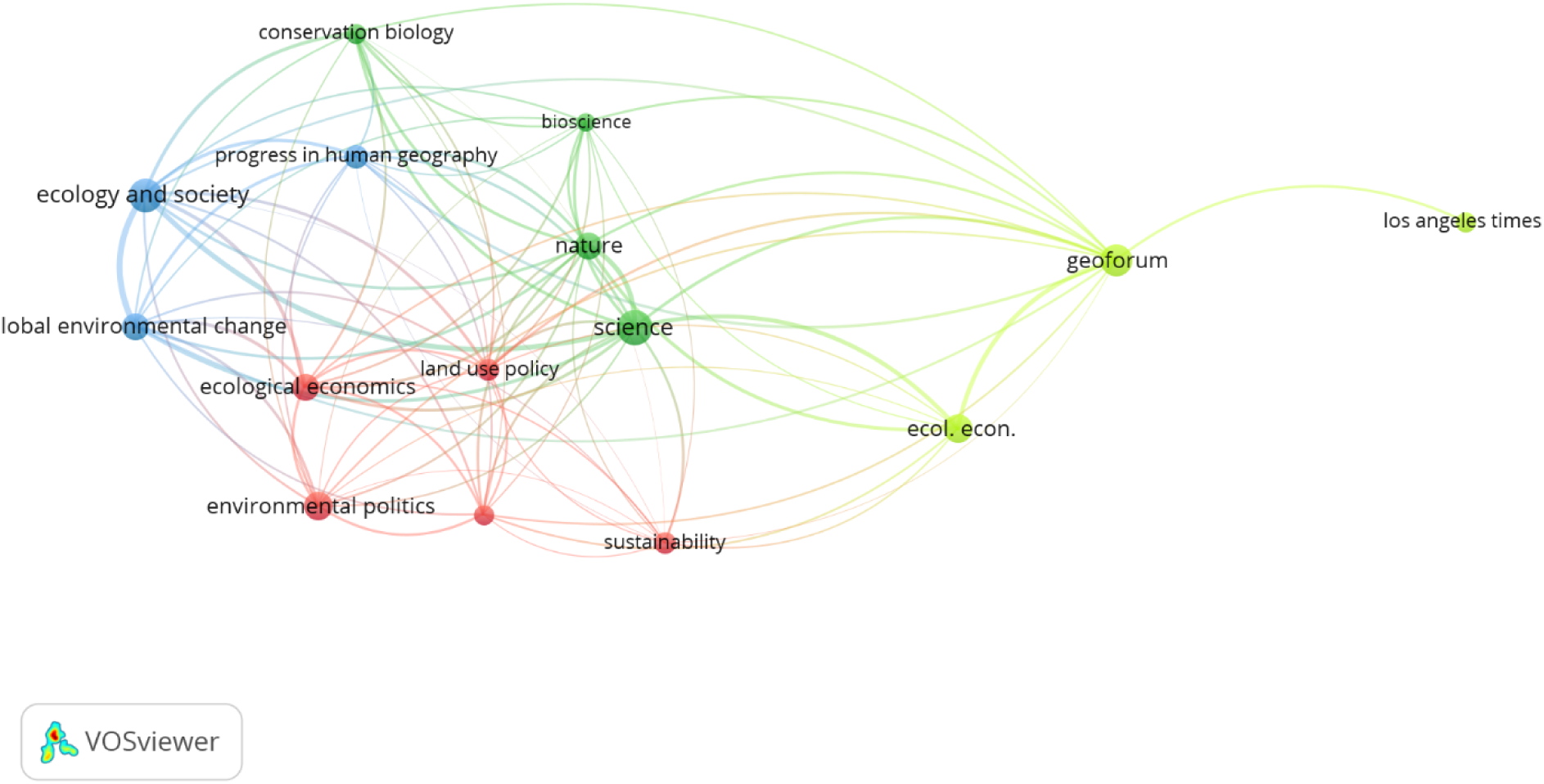
Visualization of Co-Citation Network for Fourteen Cited Sources.

**Figure 9.**
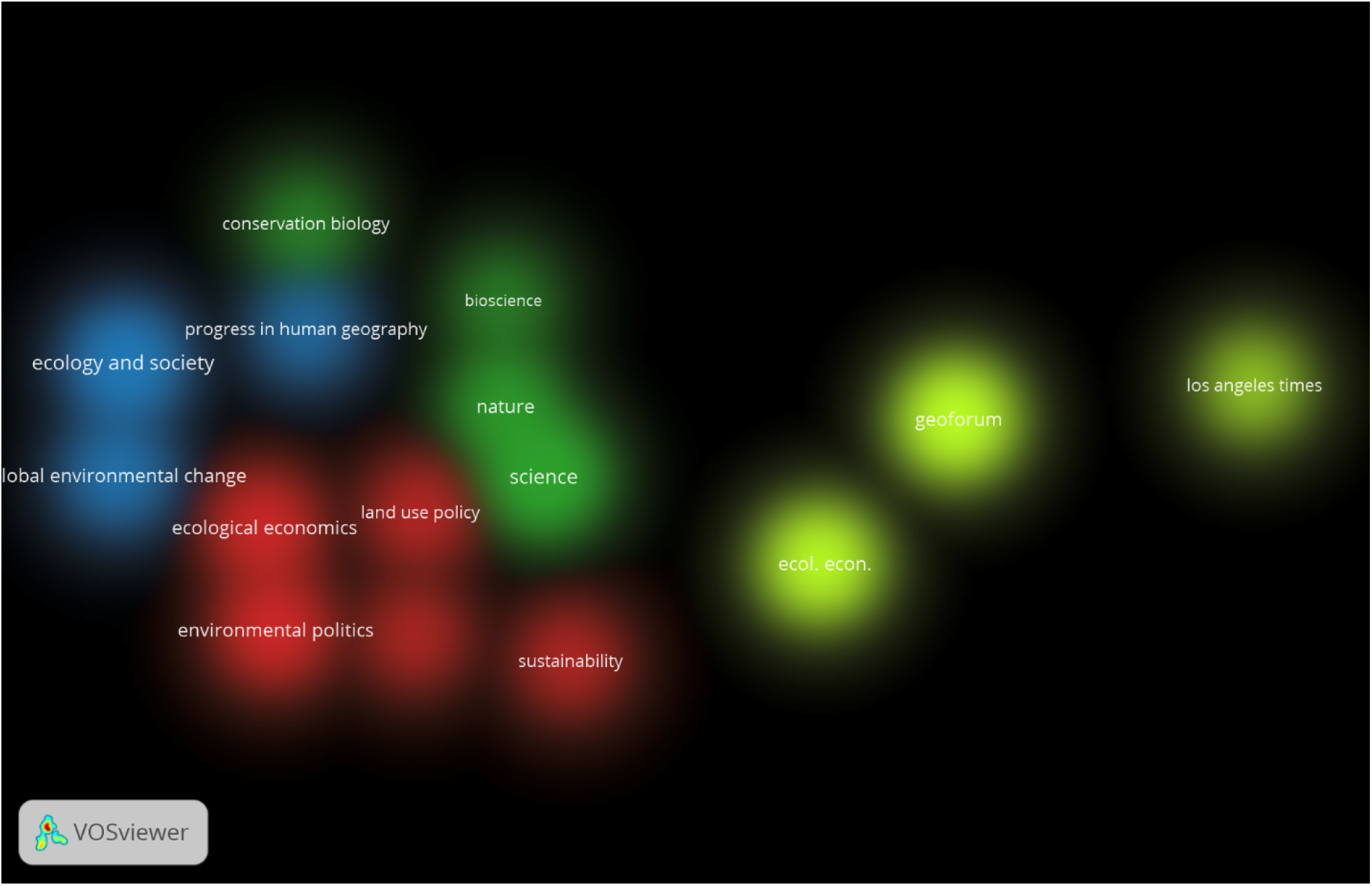
Co-Citation Density Visualization of Fourteen Cited Sources

**Table 6.** Summary of the Fourteen Most Frequently Cited Sources.

| No | source | citations | Co-citation links |
| --- | --- | --- | --- |
| 1 | Conservation biology | 55 | 385 |
| 2 | Ecol. Econ. | 111 | 421 |
| 3 | Ecological economics | 98 | 523 |
| 4 | Ecology and society | 143 | 919 |
| 5 | Energy policy | 55 | 359 |
| 6 | Environmental politics | 104 | 331 |
| 7 | Geoforum | 126 | 592 |
| 8 | Global environmental change | 96 | 723 |
| 9 | Land use policy | 67 | 386 |
| 10 | Los angeles times | 51 | 51 |
| 11 | Nature | 96 | 717 |
| 12 | Progress in human geography | 72 | 427 |
| 13 | Science | 157 | 955 |
| 14 | Sustainability | 59 | 139 |
|  | Total | 1290 | 6928 |

Figure 9 uses different colors to distinguish the various clusters of co-cited sources. Cluster 1 (red cluster) shows the intersection of *ecological economics, environmental politics, energy policy, sustainability, and land use policy journals*. These journals have over fifty citations reflecting the growing academic interest in the economic and political mechanisms underlying environmental governance and policymaking. Cluster 2 (Green Cluster) involves the *journals Nature, Science, Bioscience, and Conservation Biology*. *Science* is one of the most cited sources; the co-citation link is 955, highlighting the role of scientific research in advancing knowledge about ecological systems, biodiversity, and environmental sustainability. Cluster 3 (Blue Cluster) Notable sources include the J*ournal of Ecology and Society, Global Environmental Change Journal, and Progress in Human Geography journal.* Cluster 4 (Yellow Cluster) focused on Environmental Communication and Media. Its sources include *Geoforum journal, Ecological Economics (Ecol. Econ.) journal, and Los Angeles Times journal*.

In summary, the visualization reveals four distinct but interconnected clusters, each representing a thematic focus within environmental discourse research: scientific advancements (Green), societal impacts (Blue), policy and economics (Red), and media communication (Yellow). The centrality of *Science* and *Nature* reflects their broad influence across all clusters, while sources like *Geoforum* and *Los Angeles Times* demonstrate the critical role of communication in bridging research and public engagement. This interconnected network highlights ecological and environmental studies’ collaborative and interdisciplinary dynamics.

### Co-citation of cited authors

The co-citation analysis of cited authors serves as a complement to the co-citation analysis of references and sources. A threshold of 42 citations was established, and 16 authors met this criterion. These authors are detailed in Table 7. Among them, Folke is the most frequently cited author, with 134 citations and 1,521 co-citation links. Ostrom follows with 109 citations and 955 co-citation links. Folke et al. (2005) are particularly recognized for their contributions to global issues, including environmental sustainability, resilience, and ecosystem management, which are highly relevant across fields such as environmental studies, political science, and economics. The second most frequently cited scholar, Elinor Ostrom, is known for developing the Social-Ecological Systems (SES) Framework, a widely used approach for analyzing sustainability within social-ecological contexts. Foucault (1972) ranks third, having established a foundational methodology for discourse analysis. His work remains influential in critical theory and the social sciences, contributing to extensive citation and co-citation networks across disciplines. Fairclough, another highly cited author, introduced the three-dimensional Critical Discourse Analysis (CDA) model, which serves as a framework for examining various forms of discourse.

**Table 7.** List of the Sixteen Most Frequently Co-Cited Authors.

| No | author | citations | Co-citation links |
| --- | --- | --- | --- |
| 1 | adger w.n. | 45 | 539 |
| 2 | berkes f. | 59 | 703 |
| 3 | brown k. | 53 | 487 |
| 4 | costanza r. | 42 | 205 |
| 5 | fairclough n. | 62 | 105 |
| 6 | folke c. | 134 | 1521 |
| 7 | foucault m. | 75 | 272 |
| 8 | hajer m. | 43 | 113 |
| 9 | hajer m.a. | 46 | 107 |
| 10 | holling c.s. | 63 | 858 |
| 11 | latour b. | 60 | 339 |
| 12 | martinez-alier j. | 53 | 182 |
| 13 | ostrom e. | 109 | 955 |
| 14 | robbins p. | 47 | 235 |
| 15 | swyngedouw e. | 47 | 167 |
| 16 | walker b. | 54 | 836 |
|  | Total | 992 | 7624 |

Figure 10 presents the network visualization of the sixteen co-cited authors. VOSviewer categorizes these authors into two distinct clusters, as depicted in Figure 11. In this visualization, each node corresponds to an author, with their name serving as the label. The size of a node indicates the frequency of citations the author has received, while lines connecting nodes represent instances where two authors have been cited together in the same source. According to Figure 10, Folke appears as the most frequently cited author, represented by the largest green-colored node, which connects to the second cluster. In the density visualization (Figure 11), different colors are used to distinguish the clusters of co-cited authors, highlighting their relationships within the research field.

**Figure 10.**
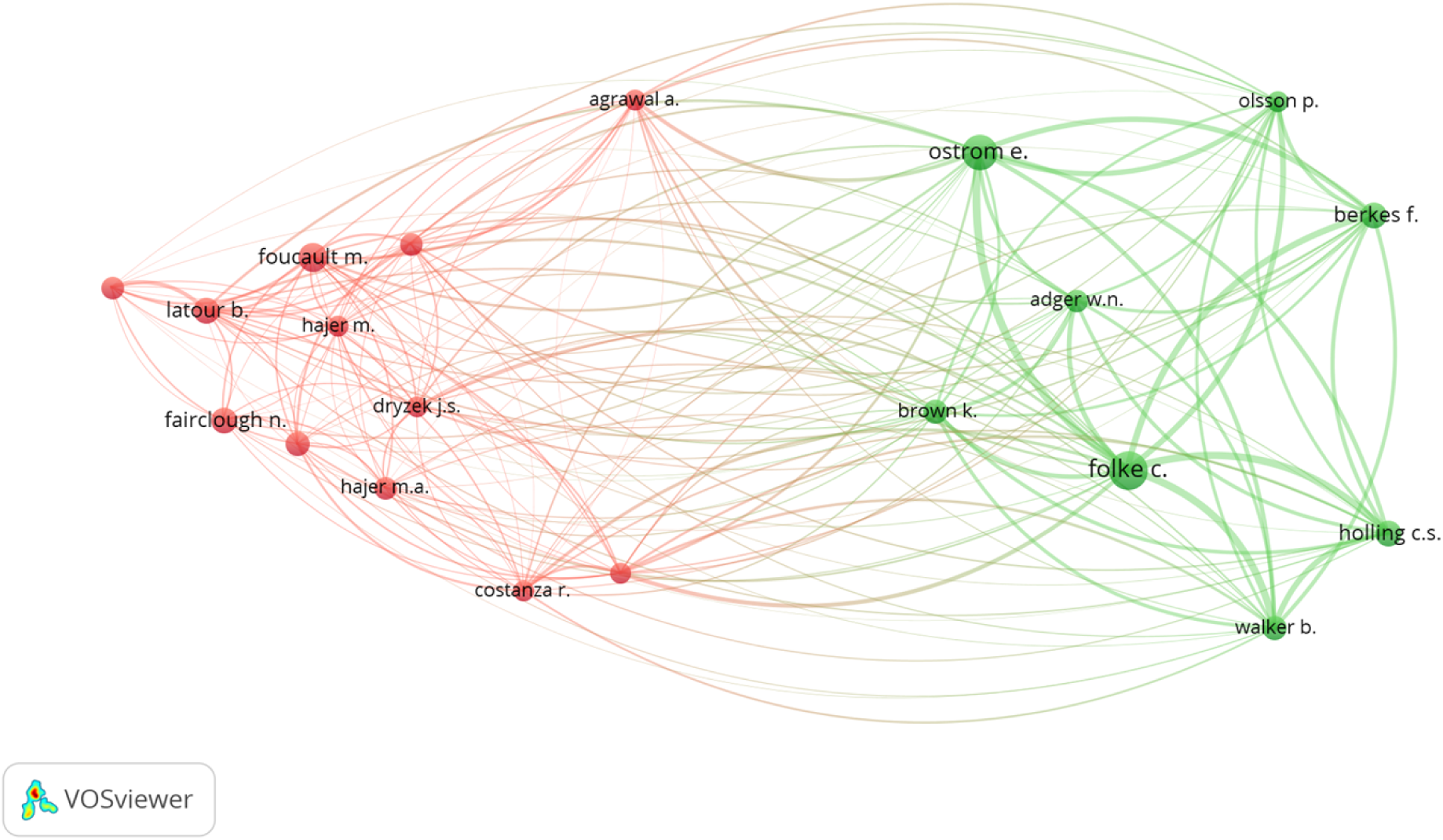
Visualization of the Citation Network for Sixteen Authors

**Figure 11.**
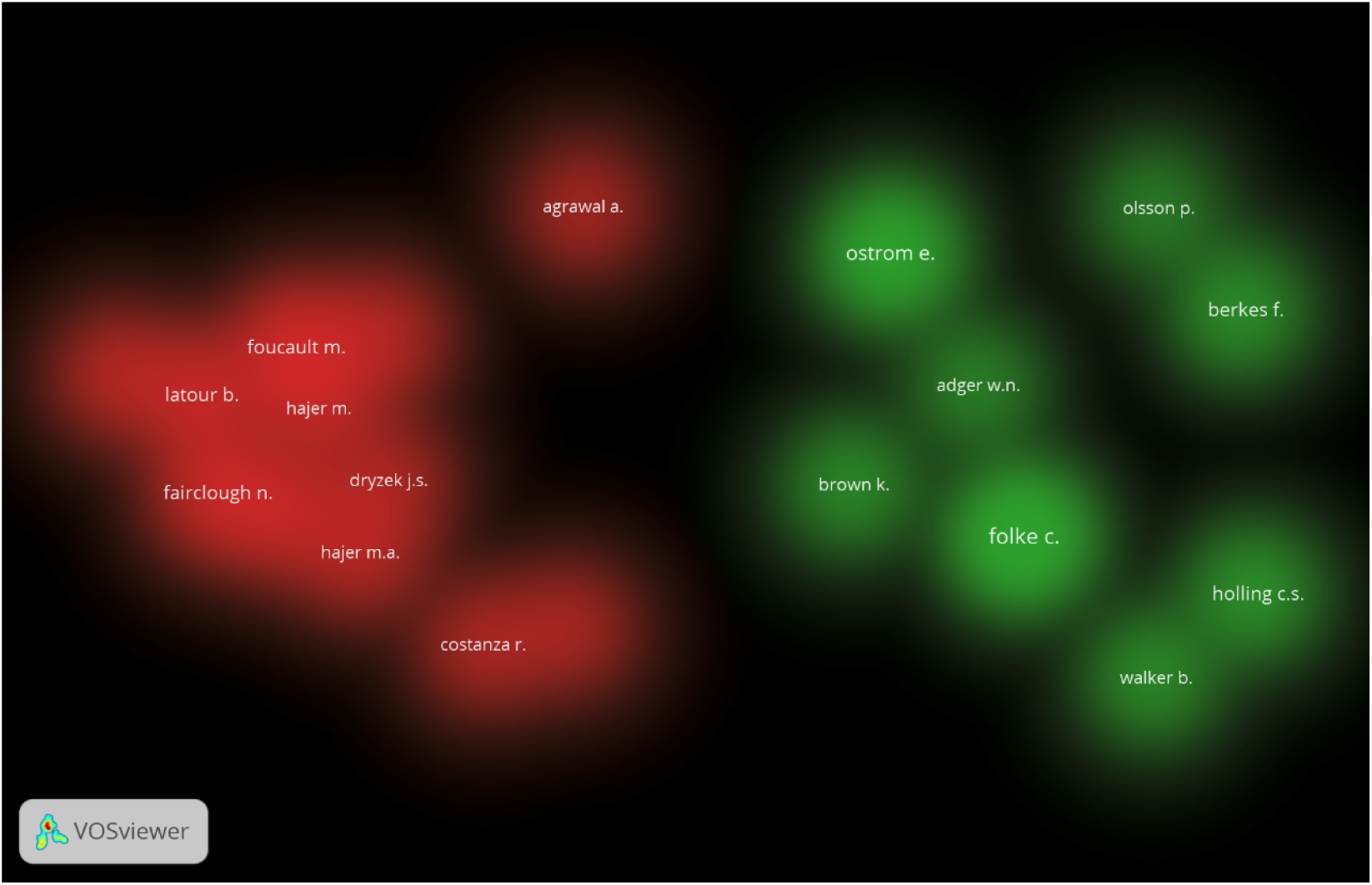
Density Visualization of Sixteen Cited Authors

Cluster 1 (Red Cluster) focuses on critical theories, environmental politics, and discourse analysis, emphasizing the interplay of power, language, and ideology in ecological discourse. Scholars such as Foucault, M., Fairclough, N., and Hajer, M. contributed to discourse analysis and environmental communication, exploring how language shapes environmental narratives and policies. Latour, Robbins, Dryzek, and Martinez-Alier. This cluster highlights ecological discourse’s linguistic, ideological, and political dimensions, incorporating critical approaches to analyze how power relations and communication influence ecological narratives. Cluster 2 (Green Cluster) focuses on ecological systems, resilience thinking, and environmental governance, emphasizing sustainability and adaptive management. Scholars include Ostrom, Holling, Walker, Folke, Berkes, Adger, and Olsson. This cluster focuses on ecological resilience, systems theory, and the governance of natural resources, highlighting interdisciplinary approaches to understanding and managing environmental challenges.

## Conclusion

To identify influential countries and sources, research themes, and the most co-cited references, sources, and authors, we used the automatic visualization tools provided by the Scopus database, the network analysis of keyword co-occurrence, and the co-citation analysis of cited references, sources, and authors. This process produced several significant findings.

First, the United States and the United Kingdom make significant contributions to ecological discourse, as reflected in their strong presence in international journals of discourse analysis. This prominence can be attributed to the well-established academic infrastructure, extensive research funding, and interdisciplinary collaborations in these countries, which foster advancements in ecological and environmental communication studies. Countries such as Germany and Australia also contribute to ecological discourse and discourse analysis from an academic perspective. Although their overall research output in this area is smaller, the papers produced in these countries are frequently cited, highlighting their influence. Researchers should identify and engage with these influential works for future studies. Additionally, countries such as Canada, Sweden, the Netherlands, China, Belgium, and Spain are increasingly contributing to research output, with China playing a significant role. China’s ecological initiatives and ideology have notably impacted global ecological awareness (Cheng, 2022).

Second, the citation analysis of sources shows that the top three most influential sources in this area are *Science, Ecology and Society, and Geoforum*. Based on the number of articles, *Sustainability, Ecology and Society, and the Ecological Economics journal* have more documents published. Although *the Journal of Environmental Policy and Planning* contributes less to this field, it receives the most citations.

Third, the network analysis of the co-occurrence of the top one hundred and eighty keywords identifies six major research themes based on the high occurrences and total link strength of the keywords: (1) Discourse Analysis, (2) Climate Change, (3) Discourse, (4) Sustainability, (5) Ecological Modernization, and (6) Critical Discourse Analysis (CDA). The synergic method of corpus and Critical Discourse Analysis (CDA) is applied extensively in ecological discourse and discourse analysis.

Fourth, it is evident that the co-citation analyses provide a historical picture of the emergence of ecological discourse analysis. The co-citation analysis of cited references shows that these are divided into six major clusters: (1) concerns political ecology and the broader sociopolitical dynamics of environmental discourse, (2) ecological systems, governance, and institutional analysis, (3) focuses on developmental discourse, human well-being, and critical discourse analysis, (4) primarily concerns discourse analysis models and political communication, (5) addresses media representation of ecological issues and environmental modernity, (6) emphasizes the philosophical and ethical dimensions of discourse. The co-citation analysis of cited sources suggests that the study of ecological discourse draws extensively on discourse analysis, among others. The frequently cited authors can be divided into two clusters: those who focus on critical theories, environmental politics, and discourse analysis, emphasizing the interplay of power, language, and ideology in ecological discourse, and those who focus on ecological systems, resilience thinking, and environmental governance, with an emphasis on sustainability and adaptive management. Combining the PRISMA guidelines and VOSviewer visualizations, this study sheds new light on the current ecological discourse and discourse analysis trends. First, the combined methodology for this study offers guidelines that show the previous literature in a staged process and improve the transparency and clarity of this bibliometric analysis; second, it helps student scholars and novice scholars identify the most productive countries, the most influential sources and documents, and the prospective authors in the field of ecological discourse studies; third, it helps scholars improve their awareness of the key topics and research trends in the field of ecological discourse and discourse analysis. The key topics and research trends are essential for scholars to follow cutting-edge research. Fourth, cited authors are clustered into Discourse Analysis and Climate Change/Sustainability. Knowing that researchers can now ground such classifications in empirical research might be beneficial. However, newer sources are becoming increasingly relevant. For instance, multimodal and critical discourse analysis in ecological discourse has attracted increasing attention recently. Therefore, the bibliometric study should continue to provide a panoramic view of the field.

This study has certain limitations. Firstly, the analysis was confined to the Scopus database, meaning that relevant studies from other sources, such as Web of Science, PubMed, or Google Scholar, were not included. As a result, some important articles indexed in other platforms may have been overlooked, potentially impacting the comprehensiveness of the findings. Secondly, the study only considered publications in English, which may have excluded valuable research in other languages.

Additionally, the study primarily relied on bibliometric tools to map co-citation networks. While these tools are useful for identifying patterns and relationships, they may not fully capture qualitative insights or the deeper interpretative aspects of ecological discourse. Future research should aim to incorporate multiple databases and integrate bibliometric analysis with qualitative methods to provide a more comprehensive perspective on the field. Despite these limitations, this study enhances understanding of key research themes and developments in ecological discourse. The findings, including frequently cited documents, widely co-cited references, and influential authors, serve as a useful resource for researchers especially those new to the field.

## Acknowledgments

This paper is part of the research achievements of the School of International Studies.

## Ethical Statement

This study, titled “Mapping Environmental Discourses: A Bibliometric and Critical Analysis of Eco-Discourses in Scopus,” presents a bibliometric analysis of eco-discourse and discourse analysis within Scopus-indexed journals. All data were sourced from publicly available secondary materials, including Scopus and bibliometric analysis tools. No human subjects or personal data were involved. The research solely analyzes existing academic publications to contribute to a better understanding of environmental discourses and their interdisciplinary connections.

## Consent to Participate

Not applicable.

## Consent to Publication

Not applicable.

## Data Availability Statement

The data used in this study were derived from the Scopus database, encompassing bibliometric metadata such as publication titles, abstracts, keywords, authors, journals, citations, and references from 2010 to 2020. The analysis was conducted using tools like VOSviewer and Scopus’ built-in analytics to ensure robust and transparent data processing. Access to the original data is governed by Scopus’ subscription and licensing policies. Processed data, including co-citation networks, keyword co-occurrence maps, and related visualizations, are available upon reasonable request to the corresponding author, subject to Scopus’ terms and conditions.

